# PinCorr: A high-pressure freezing carrier with intrinsic landmarks for cryo-correlative light and electron microscopy

**DOI:** 10.64898/2026.06.23.733922

**Authors:** Anna M. Steyer, Dietrich W. M. Walsh, Euan Pyle, Nadav Scher, Timo Zimmermann, Simone Mattei

## Abstract

Cryo-correlative light and electron microscopy methods enable targeted structural analysis of fluorescently labelled features in vitrified specimens. However, correlative workflows on high-pressure frozen samples often remain challenging due to the lack of persistent landmarks for reliable sample tracking and image registration between different microscopes. Standard high-pressure freezing carriers provide little intrinsic reference information, as the exposed sample surface is often smooth and rotationally ambiguous, complicating localisation of regions of interest across imaging platforms. Here, we introduce PinCorr, a 3-mm high-pressure freezing carrier with an integrated coordinate system formed by four asymmetrically arranged pillars with distinct geometries. These built-in landmarks remain visible after freezing and provide a stable, sample-independent reference frame for orientation and correlation between cryo-fluorescence microscopy and electron microscopy. We show that PinCorr supports fluorescence-guided cryo-volume imaging, serial lift-out for cryo-electron tomography and freeze-substitution workflows followed by room-temperature on-section correlation. PinCorr thus provides a hardware-based approach to establishing a persistent spatial reference frame in HPF-based correlative imaging workflows for thick and multicellular specimens.

## Introduction

Cryo-correlative light and electron microscopy (cryo-CLEM) has become a transformative approach for investigating the spatial and molecular organisation of biological systems across scales. Cryo-CLEM links the dynamics and molecular specificity of fluorescence microscopy with the ultrastructural and molecular context of electron microscopy under near-native, vitrified conditions. In recent years, advances in cryogenic fluorescence imaging, including high-numerical-aperture confocal microscopy and super-resolution modalities, have substantially improved the precision with which fluorescently labelled structures can be localised before downstream electron-microscopy analysis ^1 2 3^. These developments have strengthened targeted cryo-electron microscopy workflows, in which fluorescence information is used to guide focused ion beam (FIB) milling ^4^, volume imaging ^5^, or tomography towards specific regions of interest ^2^.

A central requirement in all correlative workflows is the ability to identify the same region reliably across imaging modalities that differ markedly in field of view, resolution, contrast mechanisms, and stage geometry. This step depends on fiducial landmarks or other persistent reference features that are visible in both light and electron microscopy and can therefore be used for image registration and targeting ^6–8^. For plunge-frozen specimens on EM grids, this problem is often mitigated by the grid architecture itself, by fiducial particles added to the sample ^7^, or by recognisable surface features that remain accessible during imaging. By contrast, these advantages are largely absent in samples prepared by high-pressure freezing (HPF), which is the method of choice for thicker cells and multicellular specimens that cannot be vitrified by plunge freezing ^9^.

In HPF-based workflows, the specimen is enclosed within a metal carrier and vitrified as a compact sample block ^10^. In these samples, targeting ROIs requires guidance from fluorescent signals, because after freezing, the exposed sample surface is frequently smooth and feature-poor, offering few stable landmarks for cross-modal registration ^11^. Standard carriers are also rotationally symmetric, which means that sample orientation can be lost during transfer between instruments or during loading into different holders. As a result, correlation is often slow, operator-dependent, and prone to error. These difficulties become especially limiting in workflows that require precise targeting within large sample volumes, such as cryo-volume imaging or serial liftout for *in situ* cryo-electron tomography where millimetre-scale specimens must be narrowed to micrometre-scale regions of interest ^7,12–14^. These issues also affect room-temperature correlative workflows that proceed through freeze substitution and resin embedding before TEM section imaging and tomography ^15–18^.

Existing strategies, such as the use of external fiducials ^7^, intracellular fiducials ^8^, or carrier-surface patterns ^19^, have improved specific workflows, but they do not fully solve the correlation problem for HPF specimens. Fiducials embedded in the sample cannot be detected during scanning electron microscopy (SEM) imaging of the vitreous HPF slab, whereas surface-dependent features can be lost, obscured, or altered during sample handling or surface modification, such as cryo-planing ^20^ and gas injection system (GIS) coating ^21^. An engraved FinderTOP-like approach can imprint a pattern onto the sample surface and thus support two-dimensional correlation ^19^, however the success of this method relies on reliable pattern transfer to the ice surface and does not provide a robust internal reference frame that remains intact if the sample surface is modified, obscured by ice contamination, or partially removed. Fiducials can also be milled into the ice, but this risks damaging the area of interest and can be time consuming as it requires an extra milling step ^22^.

Here, we address this problem by introducing PinCorr, a 3 mm high-pressure freezing carrier designed specifically to provide persistent, sample-independent landmarks for correlative imaging. PinCorr incorporates an intrinsic coordinate system composed of four asymmetrically arranged metal pillars with distinct geometries that extend through the sample volume to the surface of the frozen sample–carrier assembly. Because these landmarks are part of the carrier itself, they are independent of specimen morphology, and are readily visible in reflected-light cryo-fluorescence microscopy and in scanning electron microscopy. These markers provide an unambiguous orientation reference during handling and imaging. In this way, the carrier converts the otherwise featureless HPF sample block into a sample–carrier assembly with an integrated spatial reference frame. We show that this design supports robust correlation across multiple workflows, including fluorescence-guided cryo-volume imaging, serial lift-out followed by cellular cryo-electron tomography, and room-temperature correlative imaging after freeze substitution and resin embedding. By embedding the reference system directly into the HPF carrier, PinCorr provides a persistent, sample-independent coordinate system that supports cross-modal registration across multiple cryogenic and resin-based workflows for thick and multicellular specimens.

## Results

### Carrier design

The PinCorr carrier is based on a standard 3 mm B-type HPF carrier containing a 300 µm-deep well. It is designed to embed an intrinsic coordinate system into the vitrified sample–carrier assembly and to provide robust, persistent fiducials for correlative workflows. To this end, we introduced four flat-topped pillars (300 µm in height) that extend from the base of the carrier well to the carrier surface (Figure 1A). The pillars are arranged asymmetrically to define a unique orientation. Three pillars are circular (170 µm diameter), whereas the fourth is squared (200 µm side length) (Figure 1A-B).

**Figure 1.**
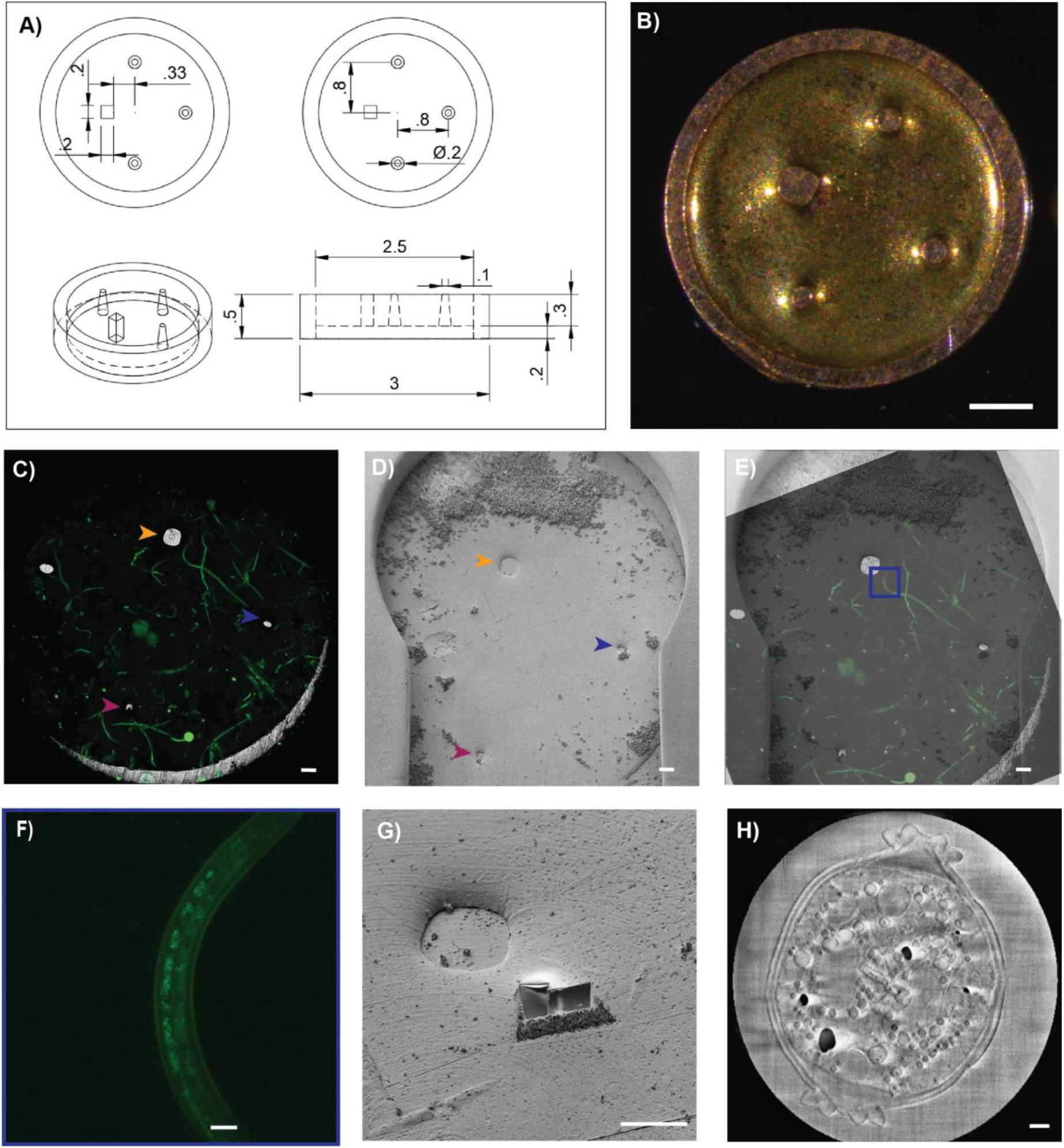
PinCorr high-pressure freezing carrier design and correlative targeting for cryo-volume imaging. **(A)** Schematic of the PinCorr showing the integrated pillar-based coordinate system. **(B)** Stereoscopic view of the carrier. Scale bar, 500 µm. (C-D) Representative high-pressure frozen *C. elegans* specimen imaged by cryo-confocal microscopy and cryo-SEM (arrow heads indicating same pillar tops): **(C)** Cryo-light microscopy overview showing the carrier landmarks in reflected light (pillar tops defining the z = 0 reference plane) overlaid with fluorescence (mNeonGreen) to localise target structures relative to the carrier coordinate system. Scale bar, 100 µm. Maximum-intensity projection of a 2×2 tiled z-stack acquired with a 10× objective (Epiplan-Apochromat 10×/0.4 NA). Scale bar, 500 µm. **(D)** Cryo-SEM surface image of the same sample acquired with secondary electrons, showing the same landmarks. Scale bar, 100 µm. **(E)** Overlay of fluorescence and SEM images after landmark-based registration by three-point alignment. **(F)** Maximum-intensity projection of the targeted fluorescence z-stack acquired with a 100× objective. Scale bar, 10 µm. **(G)** Cryo-SEM surface view of the targeted region after trench milling, exposing a cross-section through *C. elegans* at the fluorescence-defined position. Scale bar, 100 µm. **(H)** Representative slice from the cryo-volume EM dataset acquired from the exposed cross-section (secondary electron image; post-processed by denoising and stripe removal). Scale bar, 1 µm.

During assembly of the HPF sample–carrier assembly, the flat tops of the pillars contact the opposing flat face of a second, standard B-type carrier, which is used as a “lid” to close the sample sandwich (Figure S1). This geometry minimises the likelihood that sample material or cryoprotectant remains trapped between the pillar tops and the lid. After freezing and separation of the carrier sandwich, the pillar tops are exposed at the surface of the frozen sample–carrier assembly (Figure 1C). Owing to the contrast in material properties between the metal pillars and the vitrified specimen, the fiducials are readily detectable by cryo-light microscopy reflection and by cryo-scanning electron microscopy (cryo-SEM) imaging (Figure 1D–E), enabling reliable correlation between modalities.

The asymmetric arrangement and distinct pillar shapes allow unambiguous determination of sample orientation. If one fiducial is partially obscured, the remaining three are sufficient to recover orientation robustly. Pillars are positioned at least 300 µm from the carrier rim to ensure that their tops remain visible when the carrier is mounted in light- or electron-microscopy holders and are not masked by clamping elements. The inter-pillar spacing is chosen such that the maximum distance from any point within the usable sample area to the nearest fiducial does not exceed 500 µm, keeping correlation features close to regions of interest without unduly restricting sample area. Depending on the user requirements, the spacing of the pillars could be adapted.

Because the pillar-defined coordinate system is visible in both modalities, surface images can be aligned and correlated by identifying at least three corresponding fiducials and applying an affine transformation (translation, rotation, and shear). This can be readily achieved using standard microscope software in cryo-FIB workflows such as MAPs (Thermo Fisher Scientific) or Zen blue (Zeiss) as well as in open-source software such as eC-CLEM ^23^. In addition, the fiducials are large enough to be seen by eye during handling and mounting which facilitates rapid, coarse orientation of the sample–carrier assembly.

### Compatibility of the sample-carrier complex with cryo-CLEM workflows

A central design objective of PinCorr is broad compatibility with established CLEM workflows, including both cryogenic and room-temperature CLEM pipelines. We therefore evaluated whether specimens high-pressure frozen in PinCorr can be processed and imaged using fluorescence-guided cryo-volume imaging, cellular cryo-ET workflows, and room-temperature on-section transmission EM.

### Cryo-volume imaging

To assess the suitability of PinCorr for fluorescence-guided ROI targeting and cryo-volume imaging, we implemented the workflow using *C. elegans* expressing mNeonGreen in a component of the synaptonemal complex. Samples were prepared for HPF following standard procedures ^24^ using 20% Dextran or 20% BSA as a cryoprotectant/filler, by either washing the worms of the culture plates or picking them individually (wormbook, ^25^) and loading them into PinCorr carrier. Carrier were cleaned, gold coated, and glow discharged immediately before freezing and loading of the sample.

After HPF, we first imaged the vitrified sample–carrier assembly by cryo-confocal microscopy to determine the position of the specimen relative to the intrinsic, pillar-based coordinate system (Figure 1C). Light microscopy was also used to characterise the sample determining the distribution of sample (*C. elegans*) in the sample-carrier assembly as well as the distribution of the fluorescently targeted region (synaptonemal complex). This data was then leveraged to target the same area in the EM. The same carrier was then transferred to a Crossbeam 550 cryo-FIB/SEM platform (Zeiss), where a low-magnification SEM image of the carrier surface was acquired to visualise the fiducial landmarks and provide a navigation map for subsequent targeting (Figure 1 D).

To correlate the cryo-LM and cryo-SEM datasets, we registered the fluorescence and SEM overview image in Zen blue (connect module) by manually selecting at least three corresponding landmark points in both modalities and applying an affine transformation. The metal pillar tops provided high-contrast fiducials in both reflected-light cryo-LM and SEM, enabling rapid and reliable registration and fluorescence-guided identification of carrier regions containing the labelled *C. elegans* specimen (Figure 1D–E).

Acquisition of overview images of the sample surface and spatial registration between the different imaging modalities was completed in a single step by selecting the pillar tops as corresponding landmarks in both modalities. The metal pillar tops provided high-contrast fiducials in both reflected-light cryo-LM and SEM, enabling reliable registration without the need for additional preparatory steps such as fiducial milling or pattern transfer.

Alternative strategies for cross-modal registration in HPF specimens include post-freezing milling of fiducial markers into the sample surface ^22^ and the use of engraved carrier patterns such as the FinderTOP approach ^19^. The pillar-based design of PinCorr provides landmarks that are intrinsic to the carrier and therefore independent of surface transfer quality or subsequent surface modification steps. A systematic comparison of registration accuracy and workflow efficiency across these approaches was not performed in this study.

A higher-magnification confocal z-stack acquired with a 100x objective was docked into the lower magnification fluorescent atlas using common fluorescent landmarks enabling more precise localisation of the ROI within the vitrified volume (Figure 1F). Based on this fluorescence-defined ROI, we milled a vertical trench to expose an approximately 90 µm by 30 µm cross-section through the target region (Figure 1G) within the FIB. SEM imaging of the exposed face revealed a clear cross-section through the dauer larva, confirming accurate targeting. We then acquired a three-dimensional cryo-volume dataset from the targeted region (Figure 1H), demonstrating that PinCorr supports efficient fluorescence-guided targeting within large vitrified volumes.

On two different samples (from two independent freezing sessions and independent specimen) one small cube of material was acquired with approximately 10×10×4 µm (x,y,z) dimensions at a voxel size of 10×10×30 nm to give an estimate about the sample and targeting quality. Images were acquired at 2.3 kV (0.05 nA current) at 0.5 s dwell time and a line averaging of 50 using the InLens and Se2 detector. The data were postprocessed (destriped applying a variational stationary noise removal), as well as boosting the contrast and homogenizing the appearance with a local adaptive histogram normalization (CLAHE) and a gaussian filter.

### High-resolution cryo electron microscopy

Next, we performed a proof of principle experiment to validate the use of PinCorr for HPF preparation of samples to be analysed by serial lift-out and cellular cryo-ET. We tested the workflow using *C. elegans* expressing mMaple in the pharynx. The steps for sample preparation, light microscopy imaging and correlation at the cryo-SEM were essentially identical as performed for the volume imaging. The samples were used to verify if the material inside the sample-carrier assembly remained vitreous and stable enough to be thinned.

After HPF the sample–carrier assembly was imaged by cryo-confocal microscopy to determine the position of the specimen relative to the intrinsic, pillar-based coordinate system (Figure 2A), similar to the imaging in the cryo-volume workflow. Light microscopy was also used to characterise the sample determining the distribution of sample (*C. elegans*) in the sample-carrier assembly at low magnification (5/10x) as well as the distribution of the fluorescently targeted region (pharynx) at higher magnification (100x). This data was then leveraged to target the same area in the EM.

**Figure 2.**
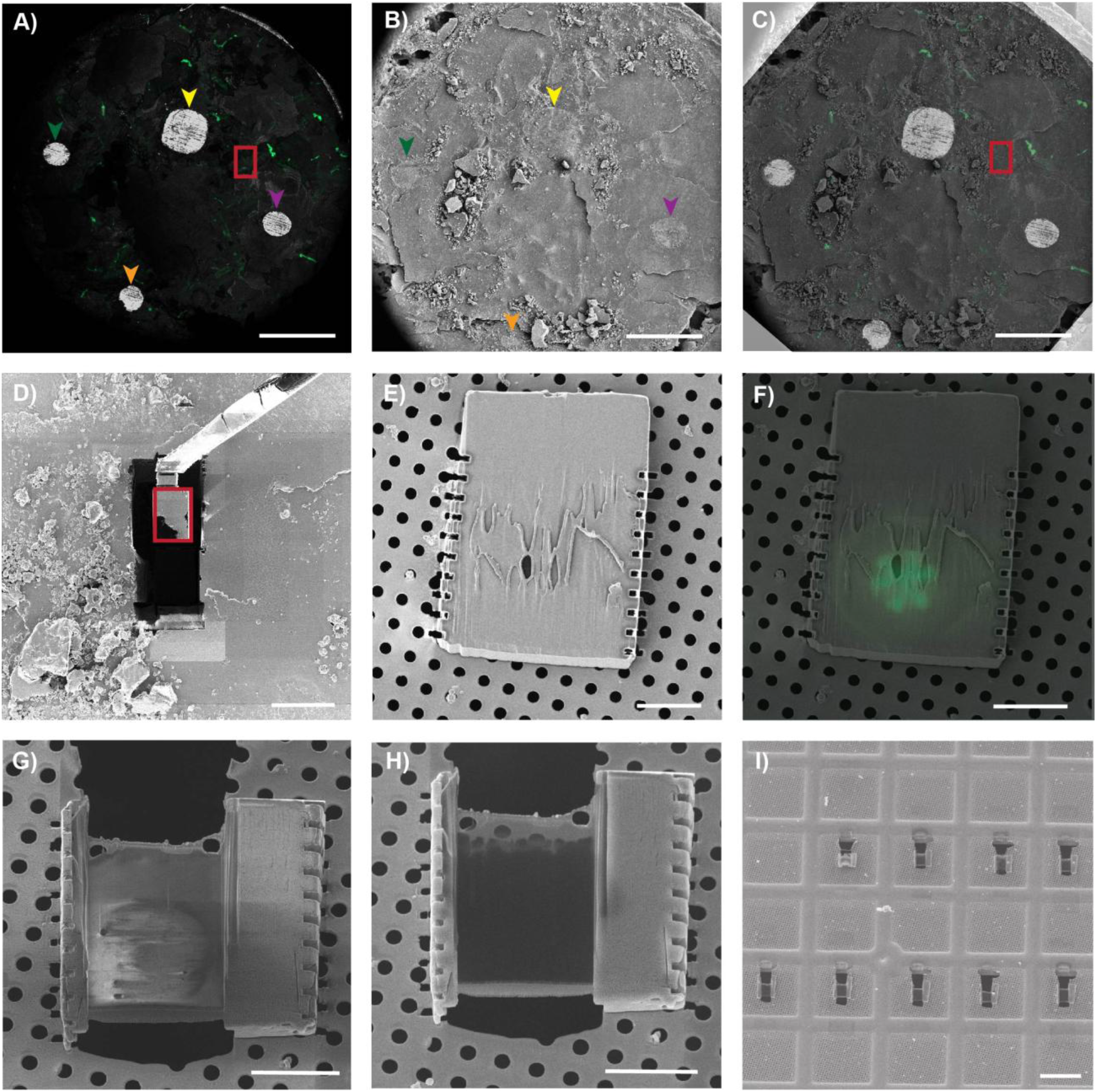
Fluorescence-guided targeting and serial lift-out from high-pressure frozen *C. elegans*. **(A)** Maximum-intensity projection of a 2 × 2 tiled cryo-confocal z-stack acquired with a 10× objective, shown in reflected light and fluorescence (mMaple, Ex 488 nm). **(B)** Cryo-SEM surface image (secondary electron signal) of the same carrier, showing the fiducial landmarks. In (A) and (B), corresponding fiducial pillars are indicated with colour-matched dashed outlines. **(C)** Overlay of the cryo-LM and cryo-SEM atlases after landmark-based registration. Scale bars in (A–C), 500 µm. **(D)** Fluorescence-guided extraction of the head region of the targeted *C. elegans* specimen by serial lift-out (SOLIST) ^14^. Scale bar, 50 µm. **(E)** A ~3 µm-thick slab transferred from the extracted block onto a TEM receiver grid. **(F)** Integrated fluorescence light microscopy (iFLM, LED 450 nm) image, overlaid with SEM image of lamella, confirming retention of the ROI (pharynx cross-section). Scale bar, 10 µm. **(G, H)** SEM images acquired at 2 kV (G) and 10 kV (H), used to assess lamella morphology and thickness. Scale bar, 10 µm. **(I)** Overview of the thinned lamella on the TEM grid. Scale bar, 20 µm.

To produce electron-transparent cellular lamellae suitable for high-resolution *in situ* cryo-electron tomography, we transferred slices of the HPF sample from the sample–carrier assembly to a receiving TEM grid. To this end, we deployed the serialized on-grid lift-in sectioning for tomography (SOLIST) method ^14^. In brief we performed correlation by selecting the same three points in the SEM surface image (Figure 2B) as were visible in the cryo-LM image (Figure 2A) using the MAPS software (Alignment wizard) to register the two imaging modalities and be able to target the chosen ROI. The obtained registered overlay (Figure 2C) allowed the identification of the ROI to be targeted by lift-out ^14^. The material surrounding the ROI was removed by cryo-FIB trench milling using an Aquilos 2 (Thermo Fisher Scientific) and an undercut was milled to isolate a freestanding block of vitrified sample. A micromanipulator needle (EasyLift, Thermo Fisher Scientific) equipped with a gold/copper adaptor was then attached to the block by ion-beam-induced redeposition, and the block was released from the bulk by a final cut (Figure 2D). The extracted block was sectioned into ~3 µm-thick slabs, which were transferred onto a receiver gold grid (Figure 2E). The integrated fluorescence light microscope (iFLM; Aquilos 2, Thermo Fisher Scientific) was used to verify slab positioning on the receiver grid and, critically, to confirm that the fluorescence-defined ROI was retained within the transferred material (Figure 2F). Lamellae were subsequently thinned and polished to the target thickness using automated routines (AutoTEM; Figure 2G–I).

The grid containing the lamellae was then loaded into the transmission electron microscope. Low-magnification overview maps were acquired to identify suitable lamella regions for tomography (Figure 3A). High-resolution tilt series were collected at multiple positions across the C. *elegans* cross-section using a dose-symmetric acquisition scheme ^26^ (Figure 3B–C). The resulting tilt series were reconstructed and denoised to generate tomograms (Figure 3D–E).

**Figure 3.**
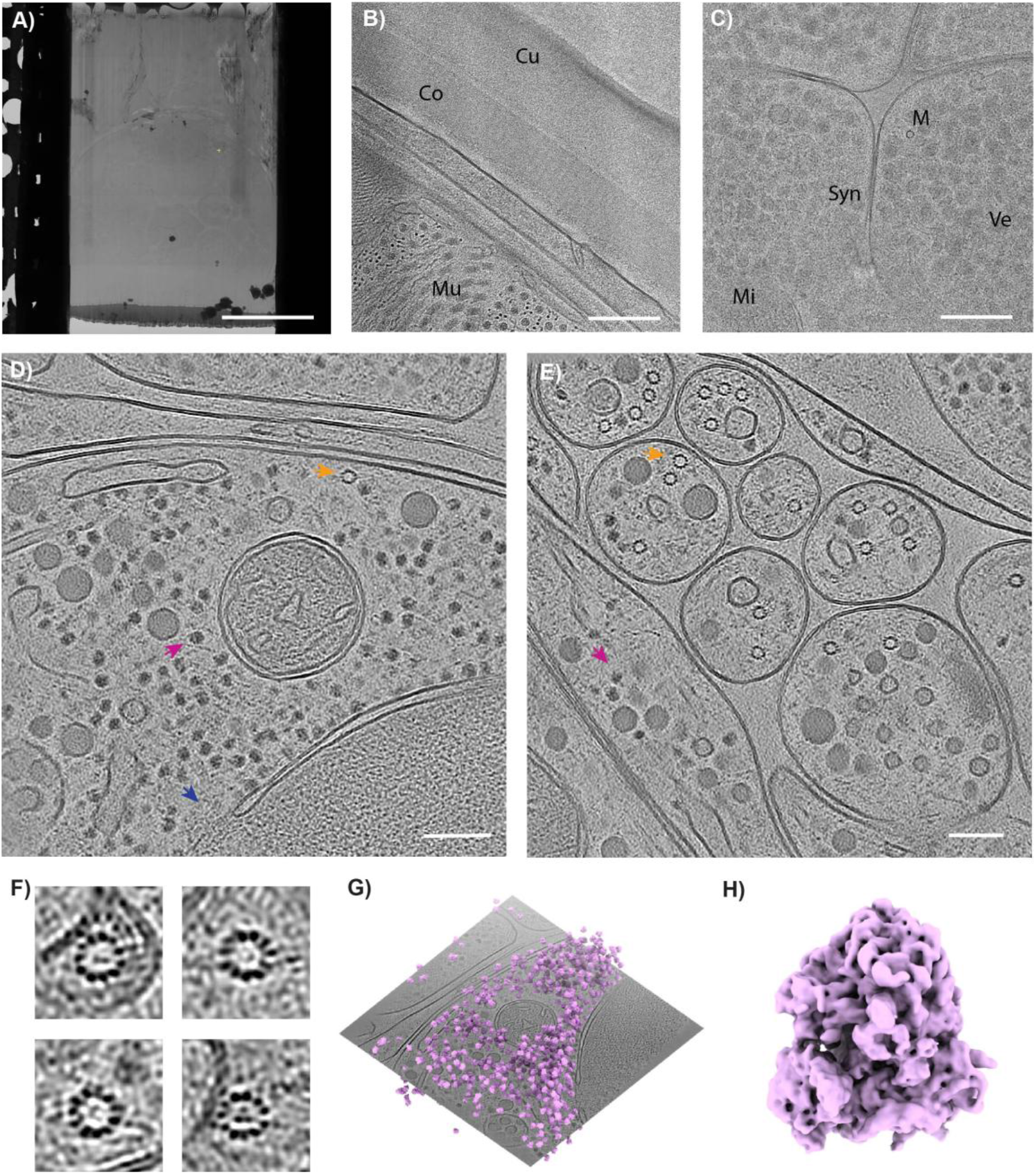
Cryo-electron tomography and subtomogram averaging from cellular lamellae. **(A)** Medium-magnification TEM overview of a lamella used for tomography. Scale bar, 10 µm. **(B, C)** Representative locally contrast-enhanced projections from a dose-symmetric tilt series (pixel size 2.405 Å). Scale bar, 200 nm. Labels: Cu, cuticle; Co, collagen fibres; Mu, muscle fibres; M, microtubule; Syn, synapse; Ve, vesicles; Mi, mitochondria. (D) Tomogram slice showing ribosomes (pink arrow), a microtubule (orange arrow), and a nuclear pore complex (blue arrow). Scale bar, 100 nm. (E) Tomogram slice from an independent lamella region showing ribosomes (pink arrow) and a microtubule (orange arrow). Scale bar, 100 nm. **(F)** Enlarged views of microtubules from the tomogram shown in (B), revealing individual protofilaments. **(G)** Tomogram slice from the lamella in (A) overlaid with the refined ribosome subtomogram average positioned at the coordinates of picked particles. **(H)** Ribosome subtomogram average at 9.1 Å resolution after refinement in M (Tegunov et al., 2021) and denoising.

Both raw tilt series and reconstructed tomograms were visually inspected during image processing and showed no obvious signs of crystalline ice. In total, 41 tomograms were generated which resolved distinct membrane leaflets, microtubule protofilaments, and individual ribosome particles (Figure 3D–F). Lamellae were close to the intended target thickness (200 nm) and had an average thickness of 219 nm (SD = 55.8 nm).

To assess whether PinCorr-derived lamellae are of sufficient quality to enable protein structure determination, we performed subtomogram averaging of ribosomes present in the cellular tomograms. Medium-magnification TEM overview of a lamella (8700x) used for tomography revealed only reflections of crystalline ice surrounding the worm tissue. We picked 7,780 ribosomes from 25 of the highest quality tomograms available and obtained a 9.1 Å reconstruction using RELION (v4.0.1) ^28,29^, followed by particle and tomogram refinement in M ^27^ (Figure 3G–H). The resulting map shows density consistent with occupancy of the A-site and elongation factor binding region (Figure 3H). The achieved resolution and map interpretability are consistent with published *in situ* ribosome subtomogram averages from serial lift-out lamellae in *C. elegans* ^13^, which reported 6.9 Å resolution using 33,350 particles. The quality of the map obtained in this study from a limited number of particles demonstrates that lamellae produced from samples vitrified by HPF using our PinCorr carrier are of sufficient quality for subtomogram averaging.

### Freeze substitution and room temperature targeting

To assess compatibility with resin-embedding workflows, we processed PinCorr high-pressure frozen *C*.*elegans* expressing mMaple in the pharynx (CHS-2::mMaple) using automatic freeze substitution following the protocol described by Ronchi et al. (2021). After the samples were embedded into a resin block, they were then sectioned and sections were observed using both light microscopy and transmission electron microscopy, targeting the same region.

Despite the different geometry and increased surface area of the PinCorr compared to a standard B-type carrier, we did not encounter major difficulties in separating the HM20 resin block from the metal carrier after embedding (Figure 4B–C). The imprints of the pillar landmarks remained visible in the resin and were used to orient and trim the block prior to sectioning (Figure 4C), providing a practical guide for reproducible positioning during ultramicrotomy.

**Figure 4.**
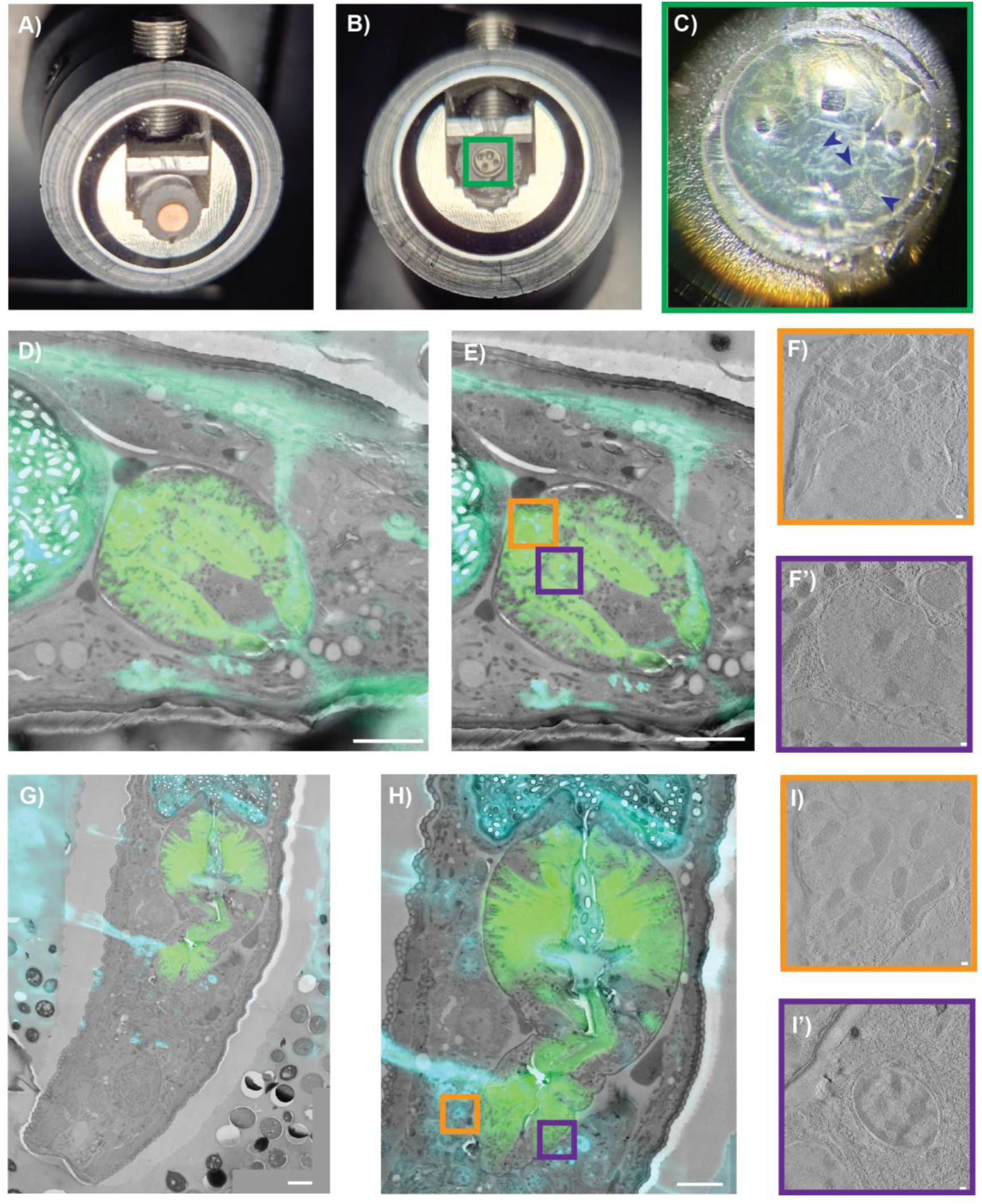
Room-temperature correlative workflows on resin embedded samples. **(A)** Polymerised HM20 resin block with the sample still enclosed in the PinCorr carrier, mounted in an ultramicrotome. **(B)** Resin block after carrier removal, exposing the embedded sample surface and the negative imprints of the pillar landmarks. **(C)** Higher-magnification view of the block surface; embedded C. elegans specimens are visible (blue arrowheads). **(D, G)** Correlation between fluorescence images of resin sections and a TEM mosaic after registration. Scale bars, 5 pm. **(E, H)** Higher-magnification views of the overlays shown in **(D. G)**. Scale bars, 5 pm. (**F, F’, I, I’**) Representative slices from room-temperature tomograms reconstructed from the regions indicated in (**E, H**). Scale bars, 100 nm.

The resulting sections showed good ultrastructural preservation, with no evidence of severe freezing artefacts (Figure 4E–F). Importantly, the retained landmark pattern also supports correlative workflows that combine freeze substitution with subsequent room-temperature imaging, including tomography of targeted structures within resin-embedded samples.

## Discussion

The main aim of this study was to address a practical bottleneck in high-pressure freezing-based correlative imaging workflows: the absence of persistent, intrinsic landmarks that support reliable sample orientation and cross-modal targeting. Standard HPF carriers generally provide little or no internal reference information after freezing, and the exposed sample surface is often smooth and feature-poor. Under these conditions, relocation of the same region between cryo-light microscopy and electron microscopy can become slow, ambiguous, and operator-dependent, particularly for large or multicellular specimens. The PinCorr carrier addresses this problem by integrating a simple asymmetric fiducial system directly into the carrier, thereby converting the frozen sample– carrier assembly into a reproducible spatial reference frame.

The data presented here support three principal conclusions. First, the pillar-based landmarks are readily detectable in reflected-light cryo-fluorescence microscopy and in cryo-SEM, enabling straightforward registration between modalities. Second, the asymmetric arrangement and mixed pillar geometry provide an unambiguous orientation cue during registration and imaging. Third, the carrier is compatible with several downstream workflows that place different demands on sample preparation and targeting, including fluorescence-guided cryo-volume imaging, serial lift-out followed by cryo-electron tomography, and freeze-substitution with room-temperature on-section CLEM. Together, these results show that the design is not tied to a single instrument configuration or a single endpoint application, but can function as a general-purpose targeting aid for HPF-based correlative workflows.

An important strength of the PinCorr design is that the fiducials are part of the carrier itself rather than added to the specimen or inferred from surface features. This makes the reference system independent of specimen morphology and less vulnerable to the variability that often complicates registration and targeting. In the proof-of-principle cryo-volume imaging experiment, correlation between cryo-LM and cryo-SEM enabled localisation of the fluorescently labelled specimen and guided trench milling to the intended region. In the serial lift-out workflow, the same principle supported selection and recovery of the fluorescence-defined region, and the resulting lamellae yielded tomograms of sufficient quality for structural analysis and subtomogram averaging. These results indicate that the carrier does not merely simplify navigation at low magnification, but is compatible with workflows that culminate in high-resolution cryo-ET readouts.

The compatibility with room-temperature resin workflows further broadens the relevance of the carrier. After freeze substitution and resin embedding, the negative imprints of the carrier landmarks remained visible in the resin block and could be used to orient trimming and sectioning. This is a useful extension of the core concept, because it shows that the reference information introduced during freezing can remain informative even after extensive downstream processing. The carrier therefore supports not only cryogenic correlative workflows, but also hybrid pipelines in which HPF is followed by resin embedding and room-temperature electron microscopy.

## Conclusion

Overall, PinCorr introduces a carrier-integrated coordinate system designed to address the absence of persistent landmarks in HPF-based correlative imaging. By embedding reference features directly into the carrier, the approach avoids dependence on variable sample surface morphology. The results presented here demonstrate compatibility with multimodal targeting across cryogenic and resin-based workflows, supporting downstream applications ranging from cryo-volume imaging to *in situ* cryo-electron tomography. PinCorr thus represents a proof-of-principle demonstration of carrier-integrated landmarks as a practical and broadly adoptable tool for correlative imaging of thick and multicellular samples.

## Acknowledgements

We acknowledge the access and services provided by the Imaging Centre at the European Molecular Biology Laboratory (EMBL IC), generously supported by the Boehringer Ingelheim Foundation. EP was funded by the Deutsche Forschungsgemeinschaft (DFG, German Research Foundation) – SPP2416 project number 525894472, and the EMBL Interdisciplinary Post-doc Program (EIPOD). We thank Martin Wohlwend for his support and effort to provide the first rounds of prototypes. We thank Julian Hennies for his support with the cryo-volume data. We would like to thank the Zimmer lab and the Köhler lab for contributing the *C. elegans* strains used in this study.

## Author contributions

Anna M. Steyer: Conceptualization, data collection, data analysis, wrote the manuscript. Dietrich W. M. Walsh: Conceptualization, wrote the manuscript. Euan Pyle: Data analysis. Nadav Scher: Data collection. Timo Zimmermann: Supervision. Simone Mattei: Supervision, wrote the manuscript. All authors contributed to data analysis and the final version of the manuscript.

## Materials and Methods

### *C. elegans* cultivation

Two different *C. elegans* strains were cultured at 20 °C and chunked every few days: (i) *C. elegans* with CHS-2::mMaple. (ii) *C. elegans* with mNeonGreen (fluorescent engineered from LanYFP, codon optimized for *C. elegans*) expressed in a component of the synaptonemal complex, these worms were cultured on nematode growth medium (NGM) agar plates with OP50 *E*.*coli* under standard conditions ^30^ at 20 °C.

### Sample preparation

Before use, the PinCorr carriers (copper) were washed in 70% ethanol and left to dry. The dry and cleaned carriers were then coated with 20 nm (30 mA) of gold using an ACE 200 (Leica Microsystems). The gold coated carriers were glow discharged right before freezing for 90 s 30 mA using the ACE 200. In parallel *C. elegans* were prepared in two ways: 1. Washed off their plates with M9 buffer and left to sediment for 5 min. The supernatant was removed and the remaining worm suspension was diluted 1:1 with 20% Dextran or 20% BSA in M9 and filled into the glow discharged carriers. 2. The worms were manually picked with a platinum rod and some *E*.*coli* paste and deposited in the carrier already containing some 20% BSA in M9. In both cases the carriers were closed with the flat side of a B-type carrier (aluminium) and either frozen using an EM ICE HPF (Leica Microsystems) or the HPM010 (Abra Fluidics).

### Light microscopy imaging

The frozen carriers were examined using a LSM900 Airyscan 2 (Zeiss) equipped with a CMS196 cryo stage (Linkam). Samples were imaged with reflected light and mNeonGreen, (Ex 488 nm Laser 4%, Emission Filter Zeiss 90 HE). Overview images were captured with a 5x objective (Epiplan-Apochromat 5x/0.2 NA) using widefield. The 2×2 tilescan and z-stack images were captured with a 10x objective (C Epiplan-Apochromat 10x/0.4 NA) with the confocal. The targeted ROI were imaged at higher magnification with a 100x objective (LD EC EPN 100/0.75 NA) acquiring z-stacks with the confocal.

### Volume electron microscopy and data processing

Samples were introduced into the focused ion beam - scanning electron microscope Crossbeam 550 (Zeiss) equipped with a cryo-stage and sample preparation system (Quorum PP3010Z). Before loading on the nitrogen gas-cooled cryo-stage, the samples were transferred in the cryo preparation chamber connected to the cryo-FIB/SEM and sputter coated with 30 mA for 40 s with platinum to dissipate charges on the surface. After introducing the sample into the main chamber of the FIB, GIS deposition was applied (opening the GIS valve for 40 s) followed by acquiring surface images with the SEM. The reflected light images collected with the cryo-confocal setup beforehand were correlated with the SEM images acquired at the cryo-FIB/SEM using Zen blue (“connect” module (Zeiss)). The LM images were already present inside a Zen connect project, which was now loaded and in which the newly acquired SEM image was saved as well. The three-point alignment tool was used with an affine transformation selecting 3 common points between the two different imaging modalities (the pins in the sample-carrier complex). To this end, the edges of the 4 pin surfaces were selected as landmarks and used as fiducials, allowing rapid registration by adapting translation, rotation, scaling, and shearing.

Based on the fluorescence the stage was then navigated to the designated sample location and a trench was produced by cryo-FIB milling to expose a cross-section of the ROI using a 15 nA current. The exposed surface was polished with 7 nA. Atlas 3D (Nanotomography, Fibics) was used to acquire the dataset using 300 pA for milling and the InLens as well as the Se2 detector (2.3 kV, 50 pA) for imaging. Since both detectors were giving slightly different results, especially regarding charging artefacts, the mixing of both signals was used to gain the best final image (mixed 50/50 within Atlas 5 (Fibics) and exported as tiff). The milling was paused when images were acquired with pixel dwell time of 0.5 µs and a line average of 20.

First, the image stacks were aligned by fixing shifts of each of the stack slices. The required translations were determined using the pre-alignment workflow implemented in the AMST2 package (https://github.com/jhennies/AMST2/) and are a combination of the ideal offset of directly adjacent slices combined with the ideal offset of a longer distance (here set to the distance of four slices). Within the AMST2 package offsets are determined by the Elastix package ^31^, on the basis of the Advanced Mattes Mutual Information score optimized by gradient descent. To remove artifacts and reduce noise, the aligned stacks were normalized (globally, histogram-based) followed by VSNR ^32–34^, local adaptive histogram normalization (CLAHE) and a gaussian filter (sigma = 1.6).

### Lift out and cryo-electron tomography

The high-pressure frozen samples were screened in the cryo-confocal and a relevant region of interest was selected based on the fluorescence, acquiring the reflected light image to visualize the fiducials as well as the fluorescent channel. The samples were transferred into an Aquilos 2 (Thermo Fisher Scientific) cryo-FIB/SEM equipped with an integrated light microscope (iFLM, Thermo Fisher Scientific) and the EasyLift (Thermo Fisher Scientific) micromanipulator needle. After imaging the sample surface, the corners/edges of the visible landmarks were used to correlate the reflected light images and select the region of interest using MAPS v3.25 (Thermo Fisher Scientific). A block of ice containing the volume of interest was lifted out following the SOLIST procedure ^14^, described briefly in the following. Sputter coating of inorganic platinum (30 mA for 30 s) was applied. A protection coating of organo-metallic platinum was applied with the gas injection system (GIS, 120 s) and to dissipate charges a sputter coating of inorganic platinum (30 mA for 30 s) was applied. A 40×30 µm block was prepared using 7 nA, milling away three sides. All edges, but most importantly the one for needle attachment, was polished using 1 nA. An undercut was performed using 3 nA leaving a block height of 25 µm to be lifted. The EasyLift needle with a copper/gold block connected was attached using multiple small rectangles (0.5 x 3 µm, 0.5 nA) to the prepared site. To release the block a side cut with 3 nA was performed. First, to make sure of the proper release, the block was moved first sideways then upwards in small steps (200 nm), then in larger steps (1 µm, 10 µm). To protect against subsequent ion beam exposure, the lifted block was re-coated with GIS platinum for 30 s before moving the block in direct contact with the recipient grid (200 mesh UltrAuFoil, Quantifoil). 3 µm slabs of material were deposited at 8° tilt onto the grid using 1 nA. The deposited lamellae were automatically thinned down to approximately 200 nm using AutoTEM v2.4 (Thermo Fisher Scientific). The following currents were used for each respective sample thickness: 1 nA to 1.5 μm, 0.5 nA to 1 μm, 0.3 nA to 0.7 μm, 0.1 nA to 0.5 μm, 50 pA to 250 nm. The final polish to the target thickness of 200 nm was done using 30 pA followed by a very last over-tilt of 0.5° (using 30 pA) to achieve a more even thickness distribution across the ice. The lamellae were sputter coated at 10 mA for 3 seconds to reduce charging during cryo-ET.

### Automatic freeze substitution (AFS)

Samples were processed using automatic freeze substitution (AFS) on an AFS-2 machine (Leica Microsystems) and the FSP pipetting robot (Leica Microsystems) as previously described ^15^. Briefly, samples were incubated for 72 h in FS media with 0.1% uranyl acetate (Serva) in glass-distilled acetone (Science Services GmbH) at −90°C then the temperature was increased to −45°C at 3.5°C/h and kept at −45°C for 5 h. FS media was exchanged three times with glass-distilled acetone, 10 min/exchange. Resin infiltration was performed by gradual increase of HM20 concentration in glass-distilled acetone (10%, 25%, 50%, 75%), 6 h for each concentration. During 50% and 75% the temperature was increased at 1.7°C/h until reaching −25°C. Then, the solution was exchanged three times with 100% HM20, 10 min/exchange. The samples were polymerized under UV light for 48 h at −25°C before the temperature was increased to 20°C at 5°C/h rate and stayed at 20°C for 6 h.

### Sectioning

Excess resin from the top and sides of the carrier was removed using a razor blade. Then the block was held above liquid nitrogen, in the vapor phase, and the carrier released using a razor blade. Blocks were trimmed using a fine razor blade, and 300 nm sections were picked on copper slot grids (AGG2500C, Agar Scientific) with a formvar film using an ultra-semi 35° diamond knife (DU3530-semi, Diatome) and Leica UC7 ultramicrotome (Leica Microsystems).

### In-section fluorescence microscopy

Section imaging was performed as previously described ^6^, with an IX81 widefield microscope (Olympus), MT20 illumination system and with a PlanApo100X 1.4NA oil immersion objective (Olympus). Images were acquired using a CCD Orca R2 camera (Hamamatsu). A 10 nm protein A gold (UMC Utrecht, CMC) solution (1:25 in distilled water) was applied to both sides of the grids as fiducials for tomographic reconstruction, before post-staining with 2% uranyl acetate in 70% methanol and Reynold’s lead citrate.

### Room temperature electron tomography

Sections were imaged with a 2100plus TEM (Jeol) at 200kV equipped with CMOS 4K or a Tecnai F30 TEM (FEI) at 300kV equipped with a OneView camera (Gatan). Single-axis tomography at regions of interest were acquired by −60° to +60° tilt series acquisition (1° increments) using SerialEM v4.1.1 ^35^, final pixel size was 1.55 nm. Tomograms were reconstructed using the IMOD ^36^ batch processing module, using gold particles tracking.

### Correlation of resin sections

Correlation of fluorescence maps to the TEM data was performed in a step-wise manner using eC-CLEM v2 plugin for Icy imaging software ^23 37^. First, the FM maps were registered to the 2300x maps using affine transformation based on sample landmarks. Then, both the transformed fluorescence map and 2300x map were registered to the 9400x maps using affine transformation based on sample landmarks. Last, both the transformed fluorescence map and 9400x map were registered to the 15500x high magnification ROIs using rigid transformation based on 10 nm protein A gold particles.

### Cryo electron tomography

Cryo-ET tilt series were collected on a Titan Krios G4 transmission electron microscope (TEM) operated at an acceleration voltage of 300 kV and equipped with a C-FEG, Falcon4 direct electron detector and a Selectris X energy filter (Thermo Fisher Scientific). Before the acquisition of the tilt series, montages of the lamella maps were acquired using 8700x magnification by selecting polygonal regions enclosing the lamellae with 30 to 35 tiles depending on the lamella size. Tilt series were collected using SerialEM v4.1.15 ^35^ with a pixel size of 2.405 Å, with a dose-symmetric scheme ^26^, starting from the pretilt angle of the lamella with respect to the grid with a total tilt range of −59 to 43, a 3° step, and a total electron dose of 150 e^−^/Å^2^ acquiring .eer frames. The energy filter slit was inserted at a width of 10 eV. A defocus of −3 to −5 µm in 0.5 µm steps was used. In total, 79 tilt series were collected using PaceTomo v1.91 ^38^.

### Data processing

Cryo-electron tomography data was processed using WarpTools (v2.0.0dev34), except where noted ^27,39^. The raw movies were converted from .eer to .tif files using RELION (v4.0.1) ^28,29^. The stage tilt angles in the .mdoc files were modified to correct for the pre-tilt of the lamella. Motion correction and contrast transfer function (CTF) estimation was carried out. Images stacks of each tilt series were generated, and manually inspected to identify low-quality images, which were deleted. Tilt series were then aligned using AreTomo2 ^40^. 3D-CTF corrected tomograms were reconstructed at 19.2 Å per pixel. The thickness of each tomogram was calculated, and the tomograms were realigned with AreTomo2 using an AlignZ value of the tomogram thickness plus 33 percent. Tomograms which were poorly aligned were either discarded or aligned using patch-tracking in IMOD ^36^. 3D-CTF corrected tomograms were reconstructed at 14.4 Å per pixel. Tomograms were denoised using cryoCARE ^41^ followed by IsoNet ^42^ for visualisation purposes only.

Ribosomes were picked using PyTOM ^43^ template matching in 3D-CTF corrected tomograms, using the structure of a ribosome from a previously obtained *C. elegans in situ* cryo-ET dataset (EMD-56322) ^44^, low pass filtered to 40 Å as the reference. The picked particles were visually inspected using ArtiaX ^45^ in each tomogram and obvious false positives were deleted. 6333 particles from 36 tomograms were extracted at 9.6 Å per pixel as 3D subtomograms. Subtomogram averaging was carried out in RELION (v4.0.1). The function relion_reconstruct from RELION was used to generate an initial reference, which was low pass filtered to 50 Å during 3D refinement. Particles were cleaned using 2D classification (K=10, T=2, no alignment), removing 326 particles, before another 3D refinement resulting in a 25 Å structure. Tomograms and particles were refined in M, using the options refine_particles and image_warp 1×1 which yielded a 20 Å structure. Tomograms were regenerated and re-denoised after M, which improved the quality of the tilt series alignment when using image_warp 1×1. PyTOM was used as before to pick ribosomes on the improved tomograms. 24219 particles were picked from 35 tomograms and extracted at 9.6 Å per pixel as 3D subtomograms before alignment in RELION. Particles were cleaned as before using 2D classification. Particles were reextracted at 4.8 Å per pixel before alignment in RELION. Duplicate particles were deleted before particles were refined in M, using the options refine_particles, image_warp 8×8, refine_volumewarp 4×4×2×35, ctf_defocus. Resolution reached 10.1 Å after M refinement. Particles were then deleted from tomograms which were visually of lower quality before M refinement as before. Particles were then extracted at 2.4 Å per pixel and refined in RELION (v5.0.1) ^46^. 3D classification (K=6, T=0.4, no alignment) was then carried out to select for high-quality particles yielding a total of 7780 particles. A last round of refinement was carried out at which point a global resolution of 9.1 Å was achieved according to Fourier shell correlation (FSC = 0.143).

**Figure S1.**
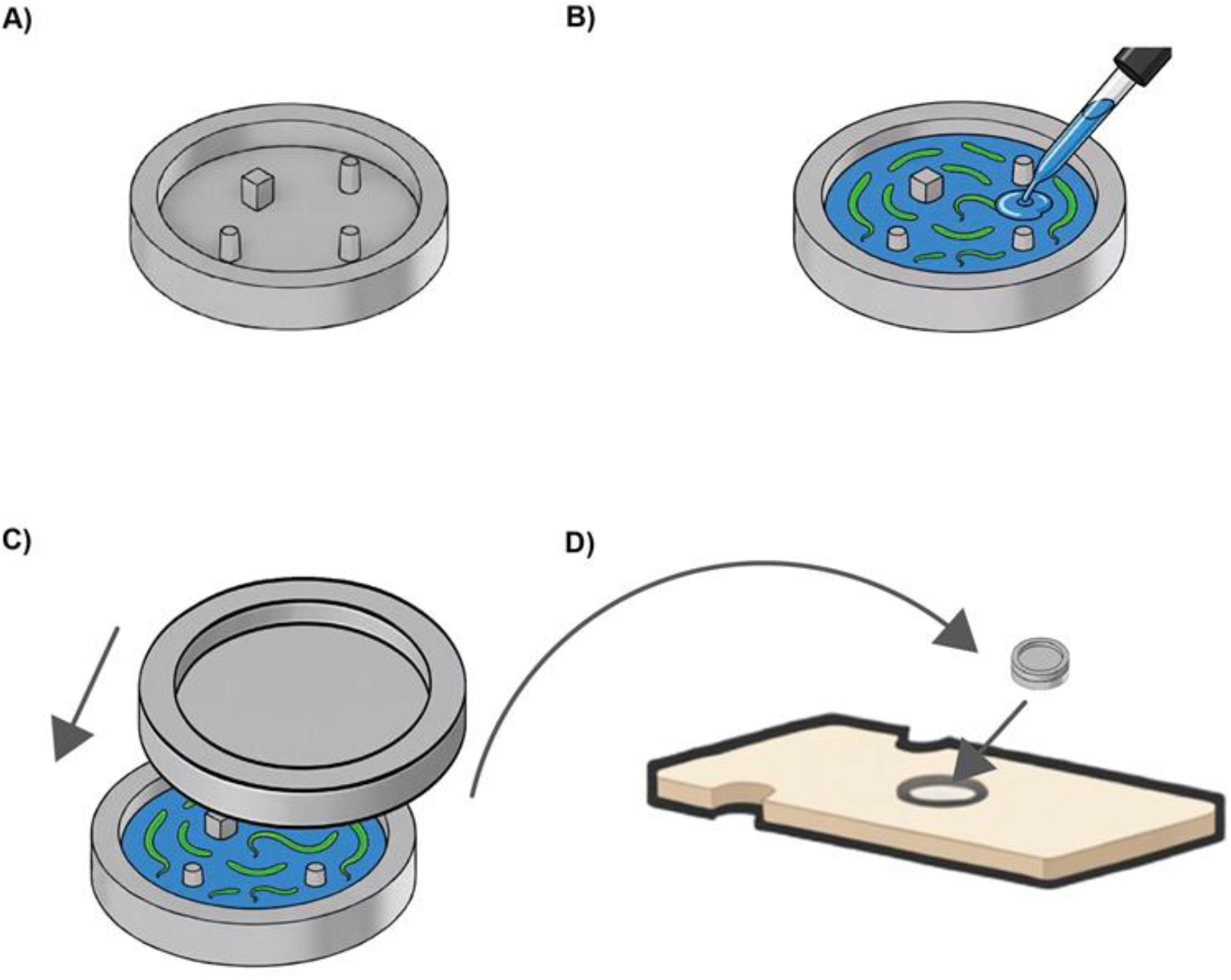
Schematic representation of sample filling and carrier sandwich assembly. **(A)** PinCorr carrier. **(B)** Filling of the PinCorr carrier with sample (*C. elegans*). **(C)** Closing of the carrier assembly (bottom: PinCorr carrier filled with sample, and top: regular B-type lid with the flat side facing down). **(D)** Positioning of closed carrier assembly in the middle plate of the high-pressure freezing cylinder before freezing.

